# Interactions of *C. elegans* β-tubulins with the microtubule inhibitor albendazole

**DOI:** 10.1101/2022.02.27.482202

**Authors:** Linda M. Pallotto, Clayton M. Dilks, Ye-Jean Park, Ryan B. Smit, Brian Lu, Chandrasekhar Gopalakrishnan, John S. Gilleard, Erik C. Andersen, Paul E. Mains

## Abstract

Parasitic nematodes are major human and agricultural pests, and benzimidazoles are amongst the most important broad spectrum anthelmintic drug class used for their control. Benzimidazole resistance is now widespread in many species of parasitic nematodes in livestock globally and an emerging concern for the sustainable control of human soil transmitted helminths. β-tubulin is the major benzimidazole target, although other genes may influence resistance. Among the six *C. elegans* β-tubulin genes, loss of *ben-1* causes resistance without other apparent defects. Here, we explored the genetics of *C. elegans* β-tubulin genes in relation to the response to the benzimidazole derivative albendazole. The most highly expressed β-tubulin isotypes, encoded by *tbb-1* and *tbb-2,* were known to be redundant with each other for viability, and their products are predicted not to bind benzimidazoles. We found that *tbb-2* mutants, and to a lesser extent *tbb-1* mutants, were hypersensitive to albendazole. The double mutant *tbb-2 ben-1* is uncoordinated and dumpy, resembling the wild type exposed to albendazole, but the *tbb-1 ben-1* double mutant did not show the same phenotype. These results suggest that *tbb-2* is a modifier of ABZ sensitivity. To better understand how BEN-1 mutates to cause benzimidazole resistance, we isolated mutants resistant to albendazole and found that 15 of 16 mutations occurred in *ben-1*. Mutations ranged from likely nulls to hypomorphs, and several corresponded to residues that cause resistance in other organisms. Null alleles of *ben-1* are albendazole-resistant and BEN-1 shows high sequence identity with tubulins from other organisms, suggesting that many amino acid changes could cause resistance. However, our results suggest that missense mutations conferring resistance are not evenly distributed across all possible conserved sites. Independent of their roles in benzimidazole resistance, *tbb-1* and *tbb-2* may have specialized functions as null mutants of *tbb-1* or *tbb-2* were cold or heat sensitive, respectively.

## INTRODUCTION

Parasitic nematodes are among the most common human pathogens and infect almost two billion people (World Health Organization 2015). Mass drug administration programs primarily use the benzimidazole (BZ) anthelmintic drug class to control infections and billions of doses have been dispensed, mainly to children (Becker *et al*. 2018). Unfortunately, previous use in livestock led to the evolution of resistance, which is now globally widespread for multiple parasitic nematode species of domestic animals (Kotze and Prichard 2016; Rose Vineer *et al*. 2020). Thus, resistance in human parasitic nematodes seems inevitable and is already emerging (Krucken *et al*. 2017; Schulz *et al*. 2018; Furtado *et al*. 2019b; Orr *et al*. 2019).

The complex life cycle of parasitic nematodes, including the requirement for obligate hosts, makes parasites difficult to study. The free-living nematode *C. elegans* has been used to study the BZ mode of action (Spence *et al*. 1982; Stasiuk *et al*. 2019; Dilks *et al*. 2020; Hahnel *et al*. 2020; Wit *et al*. 2020; Dilks *et al*. 2021). In classic work, Driscoll *et al*. (1989) screened *C. elegans* for mutants resistant to the BZ derivative benomyl and found that all 28 alleles occurred in the same gene, the β-tubulin *ben-1*. This result is consistent with the observation that BZs bind the β-tubulin subunit of microtubules and block polymerization (Friedman and Platzer 1980; Lacey and Prichard 1986; Lacey and Gill 1994; Aguayo-Ortiz *et al*. 2013). Several residues are thought to be involved in BZ binding, and these residues are mutated in β-tubulins of drug-resistant parasitic nematodes (Kwa *et al*. 1994; Kwa *et al*. 1995; Redman *et al*. 2015; Kotze and Prichard 2016; Avramenko *et al*. 2019) and confer strong resistance when introduced into *C. elegans ben-1* using genome editing (Kitchen *et al*. 2019; Dilks *et al*. 2020; Dilks *et al*. 2021).

β-tubulins are highly conserved among eukaryotes (Luduena 1998), and *ben-1* is one of six *C. elegans* β-tubulins (Hurd *et al*. 2010). The most highly and widely expressed β-tubulins in *C. elegans*, *tbb-1* and *tbb-2*, have the Y200 residue that is correlated with BZ resistance, but *ben-1* has the sensitive F200 amino acid. Although they differ in microtubule dynamics and their susceptibility to microtubule-severing enzymes, *tbb-1* and *tbb-2* act redundantly with each other for embryonic viability (Wright and Hunter 2003; Ellis *et al*. 2004; Lu *et al*. 2004; Honda *et al*. 2017). Other *C. elegans* β-tubulins, including *ben-1*, function primarily in the nervous system (Hurd 2018; Nishida *et al*. 2021).

Complete loss of *ben-1* confers BZ resistance and has no detectable growth disadvantages on or off drug under laboratory conditions (Driscoll *et al*. 1989; Hahnel *et al*. 2018; Dilks *et al*. 2020; Dilks *et al*. 2021). Indeed, many *C. elegans* wild isolates carry unique *ben-1* variants with no appreciable effects on fitness (Hahnel *et al*. 2018). In contrast, only a few point mutations appear to occur in resistant parasitic nematodes (primarily F167Y, E198A, and F200Y in the isotype-1 β-tubulin gene) although deletion of the isotype-2 β-tubulin in the small ruminant parasite *Haemonchus contortus* has also been observed in resistant isolates (Kotze and Prichard 2016; Avramenko *et al*. 2019). The limited number of mutations and the absence of clear loss-of-function mutations in the major resistance gene in parasitic nematodes, which is expected to be more frequent, imply that loss-of-function mutations would cause considerable fitness costs in the absence of drug (Wit *et al*. 2020). The reported resistant alleles likely retain function but no longer bind the BZ drugs. Although *C. elegans ben-1* is clearly the major target of BZ, other currently unknown genes can modify resistance in the field both in parasites and *C. elegans* (Zamanian *et al*. 2018; Furtado *et al*. 2019a). Mutations in the stress response, BZ uptake or metabolism modify *C. elegans* BZ sensitivity (Jones *et al*. 2015; Fontaine and Choe 2018; Matouskova *et al*. 2018; Stasiuk *et al*. 2019).

Here, we explore the role of *C. elegans ben-1* and resistance to the BZ derivative albendazole (ABZ), particularly with respect to the major β-tubulin isotypes. We found that *ben-1* is redundant with *tbb-2* as double mutants are uncoordinated (Unc) and dumpy (Dpy) in the absence of drug, phenotypes resembling wild-type animals exposed to ABZ. *tbb-2* mutants are more sensitive to ABZ than the wild type. Additionally, *tbb-1 ben-1* double mutants showed no obvious defects. These data indicate that *ben-1* and *tbb-2* are major mediators of ABZ sensitivity. As only one of the previously reported *ben-1* alleles (Driscoll *et al*. 1989) is available, we conducted a screen for ABZ resistant mutants and found that 15 out of 16 mutations occurred in *ben-1*, consistent with *ben-1* being the major target of BZs in *C. elegans*. Surprisingly, although the BEN-1 sequence is highly conserved and protein nulls are fully resistant and viable, we found that ABZ resistant missense mutations resistant to ABZ seemed to be biased toward a limited number of residues.

## MATERIALS AND METHODS

### Strains, growth conditions and ABZ treatment

Strains were maintained at 15° on NGM (nematode growth media) spread with the OP50 strain of *E. coli* as the food source (Brenner 1974). Strains are listed on Supplemental Table 1 and information about genes can be found at [WormBase](https://wormbase.org). Hatch rates were determined for complete broods of six hermaphrodites as previously described (Mains *et al*. 1990). Double mutants were made using standard genetic procedures, often aided by linked morphological markers, which were removed before analysis.

Albendazole (ABZ, Sigma #A4673) was diluted to the appropriate concentration in dimethyl sulfoxide (DMSO) so that 20-50 µL could be added to 60 cm Petri dishes containing 10 ml of NGM. This solution was quickly spread over the entire surface, and concentrations were calculated assuming uniform diffusion throughout the agar. After one day, plates were spread with OP50 bacteria, which was allowed to grow for two days at room temperature before storage of the plates at 4°. In other reports, ABZ in DMSO is often added to cooled molten agar, but we found that this procedure often forms a precipitate. Although our effective concentration may not be comparable to plates made by adding ABZ to molten agar, or to liquid culture, our results were dose dependent and reproducible even after many months of plate storage.

For measurements of larval growth, L4 hermaphrodites were transferred to the assay temperature and the next day 15-50 gravid worms were moved to fresh NGM plates without drug. The plates were incubated for approximately two hours at 25°, approximately three hours at 20°, or approximately seven hours at 11° to produce semi-synchronous broods (for the temperature-sensitive mutations *ben-2(qt1)*, animals laid embryos at the permissive temperature of 20° to bypass the temperature-sensitive period, after which they were transferred to 25°). Approximately 30-70 eggs were then transferred with a platinum wire worm pick to plates with or without ABZ. Hatching rates were near 100% in the presence or absence of drug. Plates were incubated for the specified times, during which drug-free control animals grew to the L4 or young adult stage without hatching of the next generation, which would have made it difficult to identify arrested animals. Animal lengths were measured from photographs using ImageJ software (Schneider *et al*. 2012). As we found that effects of DMSO added to NGM had no detectable effects on growth in these assays (Supplemental Figure 1A), we did not include DMSO in controls in these plate assays.

In additional to solid plate-based assays, we also performed a high-throughput phenotyping assay in response to ABZ (Dilks *et al*. 2020; Dilks *et al*. 2021). In short, a small piece of NGM agar with a starved population of individuals was placed on a new 6 cm NGM agar (NGMA) plate at 20° (Andersen *et al*. 2014). After two days, gravid adults from these plates were spot bleached to remove contamination, and the next morning, L1 larvae were placed on new 6 cm NGMA plates. These individuals were then grown for five days when a large population of L4 larval individuals were present on the plates. Five L4s were then placed on new 6 cm NGMA plates with multiple replicate populations per strain. After four days of growth, plates were bleach synchronized and the embryos were diluted to approximately one embryo per µL. 50 µL of this diluted embryo suspension was placed into each well of a 96-well plate. After these embryos hatched, these populations were then fed bacterial lysate (Garcia-Gonzalez *et al*. 2017) mixed with either ABZ in 1% DMSO or 1% DMSO alone. After 48 hours of growth, images of each well of animals were taken using an ImageXpress Nano (Molecular Devices, San Jose, CA). Images were analyzed using the easyXpress package (Nyaanga *et al*. 2021), which facilitates the measurement of individual nematode sizes from images and calculates summary statistics for sizes of populations of nematodes.

### Screen for ABZ resistance

Ethyl methanesulfonate (EMS, Sigma) mutagenesis of the wild-type reference strain N2 (HR1988) was conducted as per Brenner (1974) using 40 μM EMS for four hours at room temperature. A number of approaches and ABZ concentrations were used and screens are summarized in Supplemental Table 2. Mutagenized animals were placed on plates without drug and groups of 20-30 F1 gravid adults, which contain putative homozygous resistant F2 embryos, were picked onto plates that ranged from 1.5 to 50 μM ABZ at 20°. In some screens, mutagenized animals were picked directly to ABZ plates and gravid F1 progeny were counted after a week. Wild-type *E. coli* (AMA1004, which grows into thicker lawns than OP50) was added to all plates after one week. The extra food allowed mutations with weaker ABZ resistance more time to outgrow their non-mutant siblings. Plates were screened for movement or increased growth if we saw no movement defects (the latter yielded *sb156*). Resistant worms were transferred to fresh ABZ plates and only one strain, derived from a single animal, was retained per selection plate. As some screens were non-clonal, strains harboring identical *ben-1* DNA changes were deemed duplicates (two pairs were found, *sb146* and *sb147*; *sb143* and *sb159*). Thirteen unique *ben-1* mutations and one non-*ben-1* mutation (*sb156*) were found among 9500 haploid genomes (Supplemental Tables 1 and 2). *sb144* was outcrossed six times, and *sb151* five times, *sb163*, six times and *sb164*, one time to remove background mutations.

Additional screens took advantage of the *tbb-2 ben-1* Unc phenotype that we report here, so screens for movement on ABZ should be biased against new *ben-1* alleles. HR2038 *tbb-2(gk129)* was mutagenized as described above (Supplemental Tables 1 and 2). This screen yielded two mutants among 10,100 haploid genomes in one screen and five mutants out of 51,560 haploid genomes in another screen (Supplemental Tables 1 and 2).

### *ben-1* DNA sequencing

*ben-1* was amplified using Phusion Taq DNA polymerase (Thermo Scientific) and sequenced in four segments (Supplemental Figure 2). After amplification, DNA was extracted for Sanger sequencing from the gel using Bioneer *AccuPrep*® PCR/Gel Purification Kit. All sequencing was carried out at the University of Calgary Core DNA services.

### Statistical Analysis

Statistical analyses of agar plate-based studies were calculated in [Prism](https://www.graphpad.com) software. These worm length data were compared to controls run on the same day using a two-tailed Mann-Whitney Rank Sum test because most data sets failed normality tests. High-throughput assay data were analyzed using the R statistical environment and comparisons were made using an ANOVA with a *post hoc* Tukey HSD test. All data and scripts available on Github (https://github.com/AndersenLab/2021_Pallotto).

### Data availability statement

Strains and plasmids are available upon request. The authors affirm that all data necessary for confirming the conclusions of the article are present within the article, figures, and tables.

## RESULTS

To measure drug sensitivity, we scored larval lengths from semi-synchronized broods of embryos laid over a several hour period, followed by three days of growth on plates at 20° (unless otherwise stated). We found that 7.5 μM ABZ caused partial growth inhibition of the wild type so most of our experiments used this dose to detect either weak resistance or increased sensitivity (Supplemental Figure 1B). We also found maximal differences between control and 7.5 μM ABZ treatment of the wild type after three days post embryo laying (Supplemental Figure 1C). We present much of our data in two parts. First, we normalize each strain to the average length of the wild type run in parallel to see if mutations affect normal growth. Second, to compare relative drug sensitivities among strains, we normalize each drug treated strain to that strain’s average length on parallel, non-drug plates. A value of 1.0 signifies complete resistance. This approach should be sufficient to qualitatively divide strains into three categories: resistant, partially resistant, or sensitive. Unless otherwise stated, two-tailed Mann Whitney Rank Sum tests are used to compare data.

### *ben-1* is redundant with *tbb-2* but not *tbb-1*

*ben-1* null alleles display no mutant phenotype other than complete resistance to ABZ (Driscoll *et al*. 1989; Hahnel *et al*. 2018; Dilks *et al*. 2020; Dilks *et al*. 2021), indicating likely redundancy with other *C. elegans* β-tubulin genes. Previous work has shown that the maternal contributions of the major tubulin isotypes *tbb-1* and *tbb-2* are redundant with each other for embryonic viability (Wright and Hunter 2003; Ellis *et al*. 2004; Lu 2004). Therefore, we made double mutants of *tbb-1* or *tbb-2* null alleles with the canonical *ben-1(e1880)* mutation. Notably, *tbb-2 ben-1* double mutants showed an Unc Dpy phenotype in the absence of ABZ, but each single mutant was wild-type. These Unc Dpy phenotypes resemble the wild-type strain when it is exposed to BZ drugs (Figure 1A). This result was also recapitulated in length measurements after three days of growth (Figure 1B). The *tbb-2* and *ben-1* single mutants were respectively 0.91 and 0.93 the length of the wild-type controls run in parallel (Figure 1B). If the mutations were additive, we expect the double mutant to be 0.91 x 0.93 = 0.85 of the wild type. The observed average length was 0.67, indicating a mutual enhancement (p < 0.0001). A similar effect was not seen for *tbb-1 ben-1* double mutants where the observed value was 0.80, which was higher than the predicted expected value of 0.72. The *tbb-2 ben-1* mutant phenotype and the phenocopy in the wild type after ABZ exposure suggests that TBB-2 and BEN-1 have redundant functions in the cells that are affected by ABZ.

**Figure 1.**
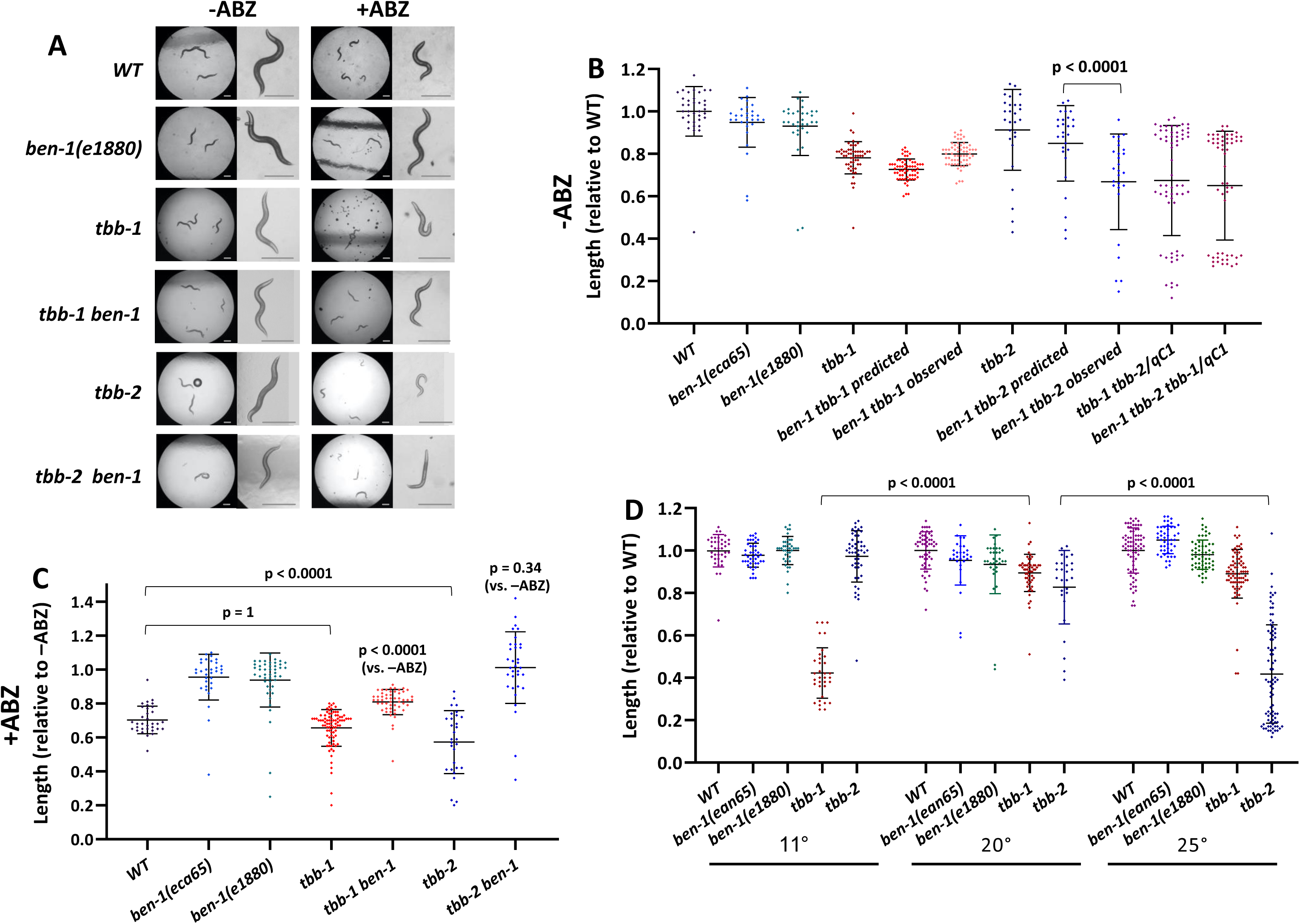
Genetic interactions of β-tubulin genes. Unless otherwise stated, experiments represent three days of growth at 20°. (A) Images of the effects of ABZ on strains at low and high magnification. *tbb-2* mutants may be hypersensitive to ABZ and *tbb-2 ben-1* double mutants grown in the absence of ABZ resemble the wild type exposed to drug. Scale bars = 250 µm. The spots in some images are crystals that sometimes form on NGM. (B) Worm lengths, in the absence of drug were normalized to the average length of the wild type. A representative wild-type sample is shown, but values of each strain were normalized to the wild type grown in parallel. The distribution of expected lengths of the double mutants of *tbb-1* and *tbb-2* with *ben-1(e1880)* were determined by multiplying each single mutant value by the average of *ben-1(e1880)*. The observed value *tbb-2 ben-1* double mutant was lower than expected. The *tbb-1 tbb-2* double mutant and *tbb-1 tbb-2 ben-1* triple mutant strains were balanced with *qC1,* which includes *dpy-19* so homozygotes are shorter than heterozygotes. The balanced strains segregate 25% arrested larvae, demonstrating that zygotic expression of *tbb-1* and *tbb-2* are redundant for viability. The addition of a *ben-1* mutation did not cause earlier arrest. (C) Effects of ABZ on tubulin mutants normalized to the average length of the same strains grown in parallel off drug. The *tbb-2* mutant appears more sensitive but was completely rescued (average near 1.0) by a mutation in *ben-1*. The *tbb-1* mutant showed normal sensitivity (but see Figure 2) and only showed partial rescue of the *ben-1* defect. (D) Although both *tbb-1* and *tbb-2* single mutants grew well at 20°, the *tbb-1* mutant was cold sensitive and the *tbb-2* mutant was heat sensitive. Growth was measured after eight days at 11°, three days at 20° and two days at 25°. Mean and standard deviations are indicated. Two-tailed Mann-Whitney Rank Sum test were used to calculate p-values.

Previous reports of redundancy between *tbb-1* and *tbb-2* were based on embryonic viability after depleting maternal stores (Wright and Hunter 2003; Ellis *et al*. 2004; Lu 2004). To determine if *ben-1* plays a role in early development, we compared the *tbb-1 tbb-2* double mutant to the *tbb-1 tbb-2 ben-1* triple mutant. As *tbb-1 tbb-2* double mutants are zygotic lethal, the double and triple mutations were maintained as balanced strains. Approximately one quarter of the larval progeny of *tbb-1 tbb-2/+* heterozygotes (presumably this one quarter are the *tbb-1 tbb-2* homozygotes) showed retarded growth, arresting once maternal stores were exhausted (Figure 1B). We observed no downward shift in the fitness of the triple mutant that included mutations in *ben-1*. It is possible that some of the triply mutant homozygotes failed to hatch and would so be missed when scoring larval lengths in Figure 1B. However, we found little to no increase in embryonic lethality in the triple mutant with *ben-1*: 3.1% of the embryos of *tbb-1 tbb-2 ben-1/+* selfed mothers failed to hatch (N=1564) comparted to 2.1% unhatched embryos for *tbb-1 tbb-2/+* (N=1356). Therefore, *ben-1* has no essential role in the embryo.

We next asked how ABZ would affect β-tubulin mutants. The *ben-1(e1880)* (amino acid change G104D) and *ben-1(ean65)* (deletion of exons 2-4) mutant strains showed near complete resistance compared to growth of the mutant off drug (Figure 1C). The *tbb-2* mutant strain was more sensitive to ABZ than the wild type (p < 0.0001), consistent with the observation that the *tbb-2 ben-1* double mutant showed mutant phenotypes off drug. Notably, *tbb-2 ben-1* grew equally well on or off ABZ (p = 0.34). By contrast, the *tbb-1* mutant showed the same sensitivity to drug as the wild type (p = 1, but see below). The *tbb-1 ben-1* double mutant was still partially sensitive to ABZ (p = <0.0001).

### High-throughput measurement of the ABZ response

We performed a high-throughput assay that is more sensitive than plate-based assays, to test how different combinations of tubulin mutations affect responses to ABZ. Populations of nematodes were grown from the L1 larval stage for 48 hours in both ABZ and DMSO (control) conditions to measure both the effects on ABZ responses and potential changes in normal growth conditions. In control conditions, we found that *ben*-*1(e1880)* mutant, the *tbb-1* deletion allele, and the double mutant grew more slowly than the wild type (Figure 2A). There was no synergistic interaction of *tbb-1* with *ben-1(e1880)* as the double mutant grew as well as the *tbb-1* single mutant (p = 0.54, we were unable to perform this assay with the *tbb-2* deletion allele or this mutation in combination with *ben-1(e1880)* because any strains harboring the *tbb-2* were too sick and slow-growing for this assay). The differences between strains in control conditions, especially the strains containing *ben-1(e1880)*, were surprising because previous studies reported that the *ben-1(e1880)* allele did not cause noticeable growth defects on plates (Driscoll *et al*. 1989). However, the highly sensitive assays used here detected a significant difference (P < 0.0001, Tukey-HSD). Conversely, the *ben-1*(*ean65*) deletion strain, which removes exons 2 to 4 created using targeted genome editing (Hahnel *et al*. 2018; Dilks *et al*. 2020), showed no changes in growth in this assay. This result suggests that the *ben-1*(*e1880*) allele likely has other mutations that affect fitness or has some neomorphic growth defects that can be revealed in liquid culture. In response to ABZ treatment, the *ben-1*(*e1880*) strain was the most resistant (Figure 2). The combination of *ben-1*(*e1880*) and the deletion of *tbb-1* was also highly resistant compared to the wild type (p < 0.0001, Tukey-HSD). The strain with the deletion of *tbb-1* alone was the most significantly ABZ-sensitive strain.

**Figure 2.**
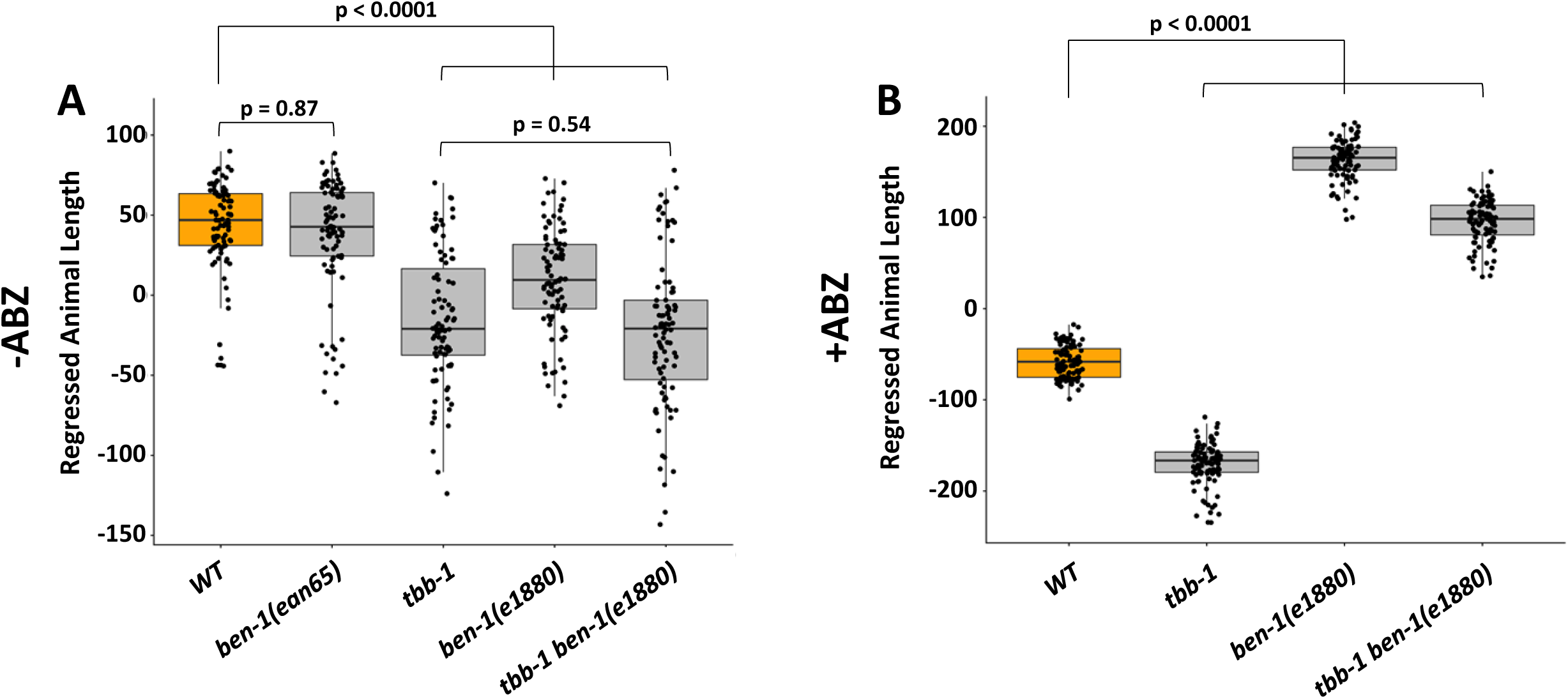
High-throughput analysis of β-tubulin mutant allele combinations. Strain names are shown on the x-axis with the allele shown under the box plot. Regressed median animal length of a population of animals is shown on the y-axis. Each point represents the median animal length calculated from a well containing approximately 50 animals. Data are shown as Tukey box plots with the median displayed as a horizontal line and the edges of the box representing the 25^th^ and 75^th^ quartiles. Whiskers are the extended 1.5 interquartile range. Tukey HSD is used to calculate significance.

### The *tbb-1* mutant is cold-sensitive and the *tbb-2* mutant is heat sensitive

Although *tbb-1* and *tbb-2* are redundant for viability, single mutants have different effects on tubulin dynamics in the early embryo and *tbb-2* also shows reduced hatching at 25° compared to *tbb-1* (Wright and Hunter 2003; Ellis *et al*. 2004; Lu *et al*. 2004; Honda *et al*. 2017). We examined temperature-dependent growth between the extremes of efficient *C. elegans* laboratory growth, 11° and 25°. We found that *tbb-1* was severely compromised at 11° (p < 0.0001 compared to 20°), and *tbb-2* showed the opposite pattern at 25° (p < 0.0001 vs. 20°, Figure 1D).

### Screens for ABZ resistance

Null alleles of *ben-1*, including large deletions, lead to BZ resistance (Driscoll *et al*. 1989; Chen *et al*. 2013; Katic and Groshans 2013; Hahnel *et al*. 2018; Dilks *et al*. 2020). Unfortunately, *ben-1(e1880)* is the only extant allele from Driscoll *et al*. (1989). To understand the range of *ben-1* mutations that can cause ABZ resistance and to possibly uncover genes other than *ben-1* that contribute to resistance, we selected for ABZ resistant mutants after mutagenesis under a variety of conditions (Supplemental Table 2, Materials and Methods). In the first screen of 9,500 haploid genomes carried out with 7.5 to 50 μM ABZ, we identified 13 independent mutants based on either movement or improved growth on drug. As discussed below, 12 had sequence changes in *ben-1*. This result yields an aggregate forward mutation rate of 1/730 *ben-1* mutations/gamete, the same frequency as found by Driscoll *et al*. (1989). This rate is higher than the average mutation rate for *C. elegans* genes of 1/2000 following standard EMS mutagenesis (Brenner 1974; Park and Horvitz 1986).

Only one mutation in the initial screen, *sb156*, lacked changes in *ben-1* (see below) and could represent an alternative target or modifier of ABZ resistance. To bias against additional mutations in *ben-1*, we carried out screens in the *tbb-2* mutant background. We reasoned that a new allele of *ben-1*, although able to grow on ABZ, would be Dpy Unc as found for the *ben-1 tbb-2* double mutant strain (Figure 1A, B). Therefore, we screened for movement, rather than either movement or growth as we did in the first screen. Some of the animals were screened at a lower dose 1.5 μM ABZ (the rest were screened at 7.5 μM) because *tbb-2* mutants are more sensitive to the drug than the wild type (Figure 1A, C). These screens yielded two mutations, but at a much lower an aggregate rate of 1/5050 per haploid genome, indicating the screen was indeed biased against frequent *ben-1* alleles. However, sequencing showed that both of these strains had *ben-1* mutations (see below). A third larger screen at 1.5 to 50 μM ABZ yielded five mutations at the low rate of 1/10,310 per haploid genome. All double mutant strains grew better than the parent *tbb-2* strain on 7.5 μM ABZ over several generations but had only marginally better movement than the parent on ABZ, especially as young larvae. These mutants have not been sequenced but are likely *ben-1* alleles for several reasons. Like *ben-1 tbb-2* double mutants, they are Unc Dpy off the drug. After outcrossing, crossovers separating resistance from the Unc Dpy phenotypes were relatively rare, indicating linkage consistent with the 2 cM map distance between *ben-1* and *tbb-2.* Therefore, although the screens in the *tbb-2* background were likely biased against *ben-1* mutations, our criterion for choosing mutants with improved movement may not have been sufficiently rigorous. Alternatively, non-*ben-1* mutations leading to resistance in the *tbb-2* background may be rare.

Several mutants had slow growth in plate assays off drug relative to the wild-type parent strain run in parallel (Figure 3A, strains are arranged in rank order of resistance found in Figure 3B). As most strains were not outcrossed after mutagenesis (Materials and Methods, Supplemental Table 2), these differences must be treated with caution. When growth of each strain on ABZ was normalized to its growth in the absence of drug, mutations varied from fully to partially resistant (Figure 3B). Although *sb156* (which lacks mutations in *ben-1*) outgrows the wild type after several generations, resistance is not seen after three days of growth at 7.5 μM ABZ and animals are Unc Dpy. Resistance was seen in the three-day growth assay at the lower dose of 1.5 μM (p = 0.005, Figure 3C, D).

**Figure 3.**
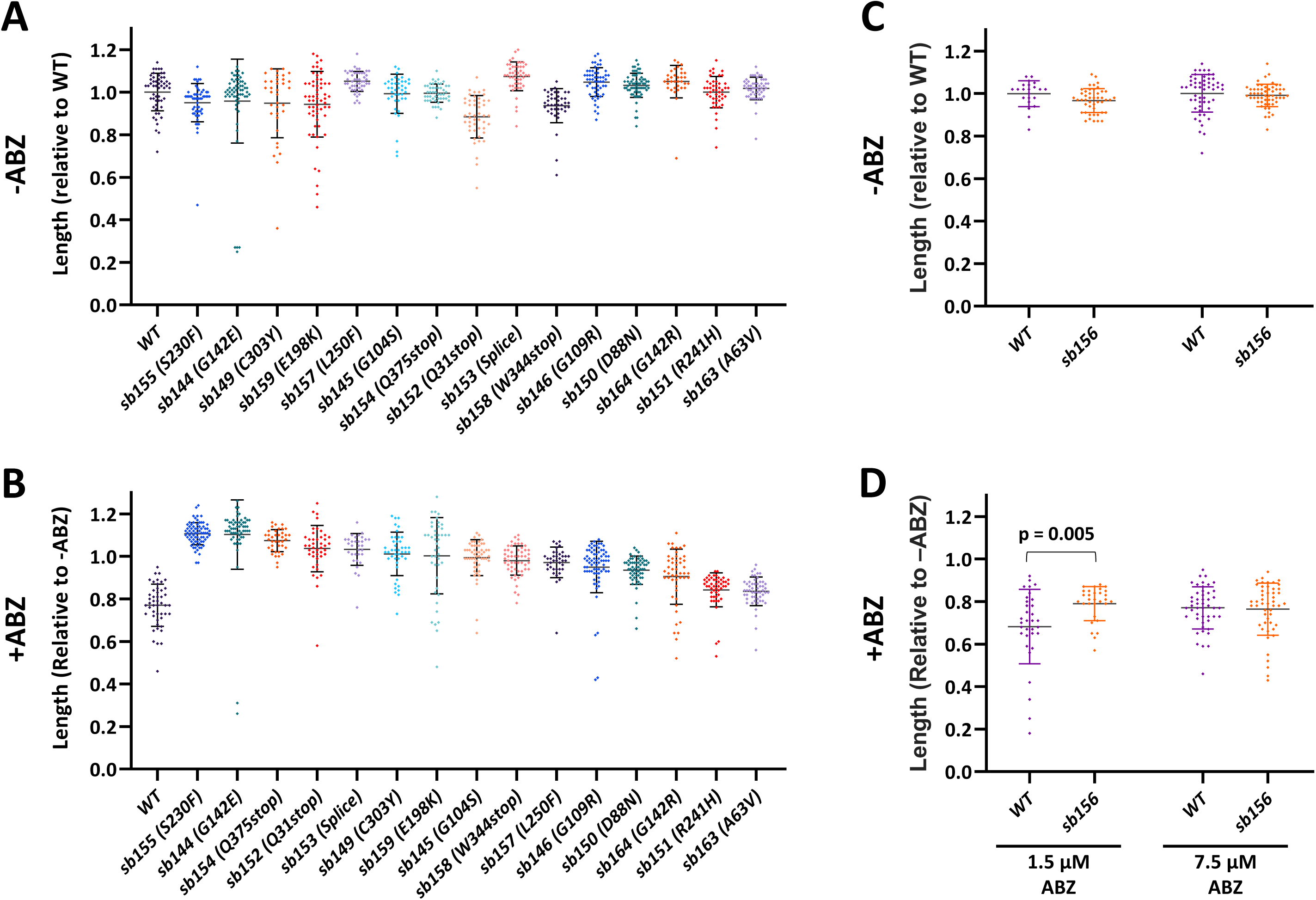
Sensitivity of mutant strains to ABZ. Strains are presented in rank order of ABZ resistance after three days of growth at 20°. (A) Growth of *ben-1* mutations relative to wild type cultured in parallel (a representative wild type is shown). (B) Growth on 7.5 μM ABZ was normalized to the average of the same strain grown in parallel in the absence of drug shown in (A). *sb151* and *sb163* showed the least resistance. (C) Growth of the wild type and *sb156*, which does not have a lesion in the *ben-1* coding region, relative to the wild type. (D) Assays run in parallel to (C) at the indicated levels of ABZ. *sb156* shows partial resistance at the lower dose. . Two-tailed Mann-Whitney Rank Sum test were used to calculate p-values. Mean and standard deviations are indicated.

### Sequence changes in ABZ resistance mutations

All mutant strains from the first two screens, except *sb156*, had sequence changes in *ben-1* (Figure 4, Supplemental Table 1). Among these mutants, several of the *ben-1* mutations are likely nulls, including nonsense alleles (*sb152*, Q31Stop; *sb158* W344Stop; *sb154* Q375Stop) as well as a splice donor mutation (*sb153,* a stop occurs after 54 intron-encoded amino acids following amino acid 157). All other mutations are in codons that encode amino acids conserved between BEN-1, TBB-2, *H. contortus* ISO-1 (the gene mutated in BZ resistant isolates of this ruminant parasite), and β-tubulins from *Drosophila*, human, and *S. cerevisiae* (Figure 4, Supplemental Figure 3 shows mutations relative to the six *C. elegans* β-tubulins). We sequenced the canonical *e1880* (G104D) allele (Driscoll *et al*. 1989) and found that it occurred in the same reside as *sb145* (G104S). Meanwhile, another pair of mutations also had different changes at a shared codon, *sb144* (G142E) and *sb164* (G142R). Two mutations have changes reported in other organisms: *sb159* E198K is found in BZ resistant fungi and parasitic nematodes and was recently found to be resistant when edited into *C. elegans ben-1* (Jung *et al*. 1992; Liu *et al*. 2014; Mohammedsalih *et al*. 2020; Dilks *et al*. 2021). The *sb151* R241H is found in benomyl-resistant *S. cerevisiae* mutants (Thomas *et al*. 1985). EMS induces GC to AT transitions and would have not induced *ben-1* mutations corresponding to the common parasite mutations F167Y, E198A, and F200Y (see Discussion).

**Figure 4.**
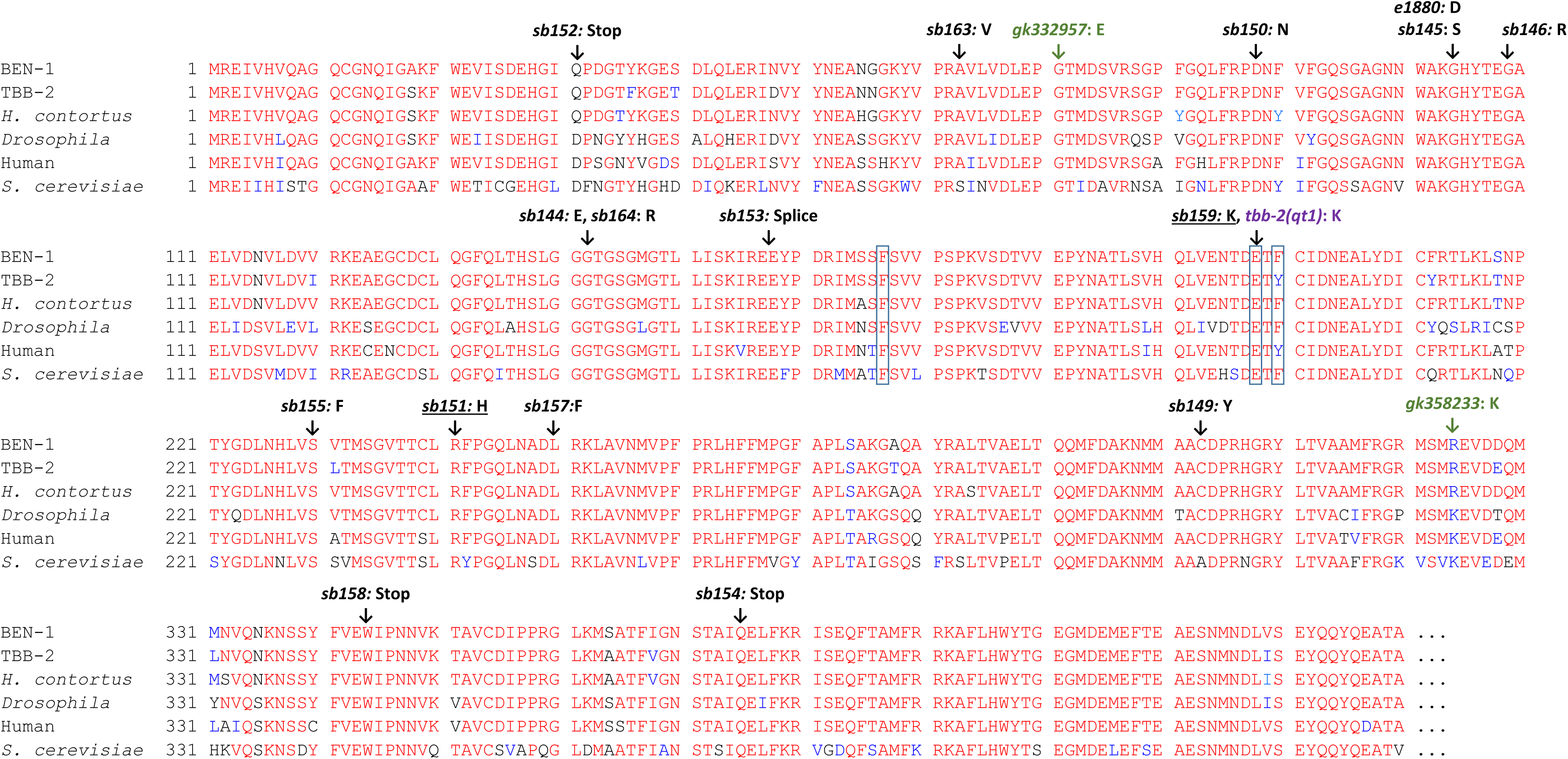
Multiple sequence alignments showing location of ABZ-resistant mutants. BEN-1 is compared to *C. elegans* TBB-2 and β-tubulins from the ruminant parasite *H. contortus*, *Drosophila,* human and *S. cerevisiae.* Locations of mutations found in our screen are indicated in black along with the canonical allele *e1880*. Boxed residues indicate the positions most frequently mutated in parasites. Underlined alleles correspond to BZ resistant mutants found in other organisms. Green represents alleles from the Million Mutant Project and purple denotes the change in *tbb-2(qt1)*. Sequences are truncated to exclude the non-conserved C-terminal regions. *H. contortus iso-1* ACS29564.1*, Drosophila* NP_651606.2, human BAD96759.1, *S. cerevisiae* NP_116616.1. For alignments to the other *C. elegans* β-tubulins see Supplemental Figure 3.

Although our data are not precise enough to correlate small changes in resistance with particular structural changes, it is notable that *sb151* (R241H) and *sb163* (A63V) had the lowest levels of resistance (Figure 3B), implying they retain some wild-type *ben-1* function. Each strain was outcrossed 5-6 times and grew well in the absence of drug, so the lack of full resistance is unlikely to come from background mutations induced by mutagen or any dominant-negative effects of the *ben-1* mutations. The alanine-to-valine of *sb163* is the most conservative change in our collection. Because *sb163* was non-Unc in combination with *tbb-2*, it likely retains some wild-type *ben-1* function. As mentioned above, the same R241H lesion seen in *sb151* is benomyl-resistant and cold-sensitive for growth in yeast. As *S. cerevisiae* has only a single β-tubulin gene, this mutation must retain wild-type function in yeast (Thomas *et al*. 1985). In analogy with the yeast R241H mutations, we tested *sb151* for cold sensitivity. Like *ben-1* null alleles, *sb151* had little affect growth at 11°, 20°, or 25° in the absence of drug (Figure 5A). If *sb151* compromises normal *ben-1* function more at lower temperatures, it should be more resistant and we found a slight increase of resistance at 11° (p < 0.0001 vs. 25°, Figure 5B).

**Figure 5.**
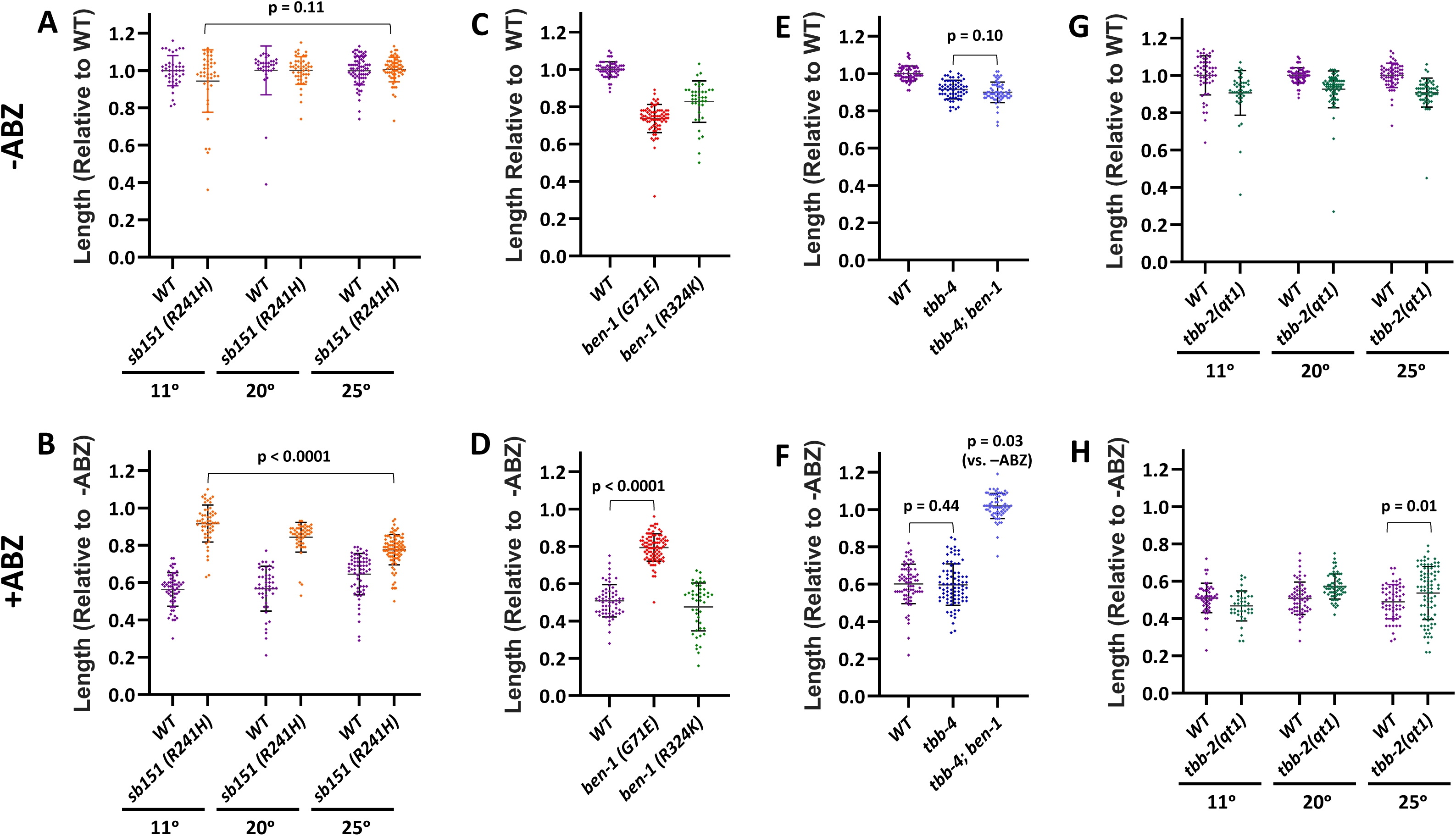
Phenotypes of selected mutations in *ben-1* and other tubulin genes. Animals were measured after eight days at 11°, three days at 20° and two days at 25°. Unless otherwise indicated, experiments were performed at 20°. Upper panels (A, C, E, G) were grown in the absence of ABZ and were normalized to the wild type run in parallel. Lower panels (B, D, F, H) are the corresponding experiments grown on 7.5 μM ABZ and are normalized to the strain grown in parallel in the absence of drug. (A, B) Although the corresponding *S. cerevisiae* is cold-sensitive for growth, *sb151* was not. *sb151* did cause a slight reduction in *ben-1* function at 11° as indicated by better growth on ABZ at the lower temperature. (C, D) The *ben-1* G71E mutation (*gk332957*) from the Million Mutant Project showed partial resistance to ABZ but R324K (*gk358233*) did not, indicating that the latter mutation does not compromise function. (E, F) Loss of *tbb-4*, which includes the sensitive F200 residue, did not slow growth in double mutants with *ben-1* in the absence of ABZ nor did it alter drug sensitivity. (G, H) The temperature-sensitive mutation *tbb-2(qt1)* showed slight resistance at 25°. Two-tailed Mann-Whitney Rank Sum test were used to calculate p-values. Mean and standard deviations are indicated.

### ABZ resistance in other β-tubulin mutant genes

The *C. elegans* genome encodes three other β-tubulin genes in addition to *ben-1, tbb-1*, and *tbb-2*. We tested two *ben-1* mutations generated by the Million Mutant Project (Thompson *et al*. 2013) that have amino acid changes found in other *C. elegans* wild-type β-tubulin genes and so might be considered conservative substitutions that retain function (Figure 4). These mutants were identified after random mutagenesis without subsequent selection for ABZ resistance. The *ben-1(gk332957)* G71E change is shared with the divergent β-tubulin TBB-6 (Supplemental Figure 3). This mutation nevertheless compromises but does not eliminate wild-type *ben-1* function as it shows partial resistance (p < 0.0001 vs. the wild type, Figure 5C, D). The *gk358233* R324K allele is likely a permissible change as it is found in fly, human, and yeast β-tubulins as well as TBB-4, TBB-6, and MEC-7 (Figure 4, Supplemental Figure 3). As expected for a functional protein, this allele was still sensitive to ABZ (Figure 5C, D). Both of the Million Mutant Project alleles showed compromised growth off drug, but the strains were not outcrossed.

The *tbb-4*, *tbb-6*, and *mec-7* genes each encode the F200 residue that correlates with BZ sensitivity, but mutations in these genes are not found in wild ABZ resistant strains (Hahnel *et al*. 2018). As *tbb-4* is expressed in some of the same neurons as *ben-1* (Hao *et al*. 2011; Nishida *et al*. 2021), we asked if loss of both *tbb-4* and *ben-1* would alter growth, similar to the *tbb-2 ben-1* double mutant. However, we found no changes (p = 0.10) off of drug and the strain was still resistant when exposed to ABZ (Figure 5E, F).

Although TBB-2 is predicted not to bind ABZ as it is Y200, the *qt1* allele has been implicated in benomyl resistance (Wright and Hunter 2003). This *tbb-2* mutation encodes the E198K change that is ABZ resistant when edited into *ben-1* (Dilks *et al*. 2021) and is present in the *ben-1(sb159)* allele from our screen. It is also found in BZ resistant parasitic nematodes and fungi (Jung *et al*. 1992; Liu *et al*. 2014; Mohammedsalih *et al*. 2020). Wright and Hunter (2003) found that *tbb-2(qt1)* prevented embryonic spindle orientation defects caused by benomyl microtubule depolymerization at the first embryonic cleavage, particularly at higher temperatures. We found a small increase in resistance in *tbb-2(qt1)* in our growth assays at the restrictive temperature of 25° (p = 0.01, Figure 5E, F).

## DISCUSSION

The World Health Organization includes ABZ on its list of [100 Essential Medicines](https://list.essentialmeds.org/medicines/57). Billions of ABZ doses have been administered for treatment of parasitic nematodes, mainly to children (World Health Organization 2017). Previous widespread use of BZ in agriculture caused the evolution of resistance, often rendering BZ drugs ineffective for a number of livestock parasitic nematode species. Resistance amongst human helminths seems highly likely and concerns are growing about its emergence (Moser *et al*. 2017). β-tubulins are the major target of BZ drugs in both fungi (Thomas *et al*. 1985; Jung *et al*. 1992; Liu *et al*. 2014) and nematodes (Driscoll *et al*. 1989; Kwa *et al*. 1995; Wit *et al*. 2020). To better understand the genetics of BZ resistance, we used *C. elegans* as a model. The β-tubulin *ben-1* gene was known to be the major target of the BZ class of drugs (Driscoll *et al*. 1989; Hahnel *et al*. 2018). We explored genetic interactions of *ben-1* and ABZ with the major β-tubulin isotypes, *tbb-1* and *tbb-2,* and conducted forward genetic screens to examine the types of mutations that lead to ABZ resistance.

### Interaction of *ben-1* with other β-tubulin genes

Assigning paralogous functions among β-tubulins within an organism, or inferring homology by descent of β-tubulins between organisms, is problematic because of their slow rate of evolution. The exception to tubulin conservation occurs in the C-terminus, which shows little similarity between tubulin paralogs within a species or between tubulins from different species. A few specializations have been ascribed to these regions (Hurd 2018). Although mutations in *C. elegans ben-1* and *iso-1* of the ruminant parasite *H. contortus* both confer BZ resistance, which might imply homology, levels and cellular patterns of expression may be more critical to define shared functions than primary sequence (Saunders *et al*. 2013).

To better understand BZ resistance, we sought to clarify the functional relationships between *ben-1* and the major β-tubulin isotypes *tbb-1* and *tbb-2.* Unlike *ben-1,* neither *tbb-1* and *tbb-2* are predicted to bind BZ as they encode Y200 rather than the sensitive F200 residue. The *tbb-1* and *tbb-2* genes act redundantly with each other for viability, both maternally (Wright and Hunter 2003; Ellis *et al*. 2004; Lu *et al*. 2004) and zygotically (Figure 1). Of these two genes, we found that *tbb-2* shows greater functional overlap with *ben-1.* In the absence of ABZ, *tbb-2* and *ben-1* are redundant for movement, body morphology, and growth (Figure 1). For these phenotypes, the *tbb-2 ben-1* double mutant resembles the wild type exposed to ABZ. A simple model is that TBB-2 and BEN-1 are expressed in the cells responsible for the ABZ- induced phenotypes. Consistent with this observation, *tbb-2* mutants had a greater increase in ABZ sensitivity relative to the wild type than did loss of *tbb-1* (Figures 1 and 2). The phenotypic similarities between *tbb-2* mutants and *ben-1* mutants could be caused by shared isotype-specific functions. Another possibility is that the overall higher levels of *tbb-2* expression relative to *tbb-1* (Nishida *et al*. 2021) could be important. If the stronger interactions that we observed in *tbb-2* double mutants are simply a matter of higher overall β-tubulin levels, it might be possible to increase BZ toxicity in parasites with subclinical doses of microtubule inhibitors that target all microtubules, rather than only those microtubules that include β-tubulin isotypes with F200.

### *tbb-1* and *tbb-2* have non-overlapping functions

*tbb-1* and *tbb-2* are redundant for viability although single mutants of *tbb-1* and *tbb-2* have only subtle effects on tubulin dynamics and modest effects on hatching rates (Wright and Hunter 2003; Ellis *et al*. 2004; Lu *et al*. 2004; Honda *et al*. 2017). If each member of a redundant gene pair efficiently provides the same functions, selection might not act to preserve both members of the pair (Nowak *et al*. 1997). However, if the members of the gene pair also have non-overlapping essential functions, selection will retain both copies. Indeed, *tbb-1* and *tbb-2* may be specialized for growth at different temperatures, as *tbb-1* mutants grew poorly at 11° and *tbb-2* mutants had compromised growth at 25° (Figure 1). This range matches the substrate temperatures of *C. elegans* collected from the Hawaiian Islands (4° to 23°) (Crombie *et al*. 2019; Crombie *et al*. 2021).

### Only certain BEN-1 residues might mutate to cause ABZ resistance

As only one allele from the Driscoll *et al*. (1989) screen for benomyl resistance is available (*e1880*), we conducted forward genetic screens to explore the types of mutations that can confer ABZ resistance. Consistent with the idea that *ben-1* loss leads to ABZ resistance (Driscoll *et al*. 1989; Hahnel *et al*. 2018; Dilks *et al*. 2020; Dilks *et al*. 2021), we found a number of nonsense alleles (*sb152*, *sb158*, *sb154*) and a splice donor mutation (*sb153*) among the 16 alleles we sequenced. These mutations are likely protein nulls and are distributed throughout the gene (Figure 4). One might suspect that the high conservation of β-tubulins would imply that most *ben-1* residues are critical for function *a priori* and so our screen could have identified missense mutations in a large proportion of the conserved sites. Outside the non-conserved C-terminus, *ben-1* shows 77% and 97% identity with β-tubulins from the yeast *S. cerevisiae* and the parasitic nematode *H. contortus*, respectively. Comparisons to other *Caenorhabditis* species also indicate that *ben-1* evolution is highly constrained (Hahnel *et al*. 2018). However, of the 12 missense mutations (we include the canonical allele *e1880* in this total), seven have what might be considered unusual properties.

Several lines of evidence indicate that relatively few *ben-1* missense mutations can lead to sufficient loss of activity to confer resistance. For example, we found two mutations that correspond to BZ resistant mutations in other parasites and fungi. In those organisms, the genes are essential, so mutations likely represent specific changes that block drug binding and retain sufficient function to compete in the wild in the absence of drug. Such mutations in the non-essential *ben-1* gene should be rare compared to those mutations that cause loss of function, if most *ben-1* missense mutations were to confer resistance. Fifteen mutations leading to resistance have been reported in parasitic nematodes and fungi. However, only three can be created in one step in *ben-1* as GC to AT transitions, which accounts for 90% of EMS-induced *C. elegans* mutations (Thompson *et al*. 2013) (Supplemental Table 1) and these mutations do not include the F167Y, E198A, and F200Y commonly found in resistant parasites. EMS can induce H6Y found in *Aspergillus nidulans,* E198K found in *A. nidulans*, *Gibberella zeae*, and *H. contortus*, and R241H found in *S. cerevisiae* (Thomas *et al*. 1985; Jung *et al*. 1992; Liu *et al*. 2014; Hahnel *et al*. 2018; Mohammedsalih *et al*. 2020). We found two of these three mutations, *sb159* (E198K) and *sb151* (R241H). *sb163* also appears to be another mutation that retains function as it was selected to confer movement in the *tbb-2* background. Another three ABZ resistant missense mutations have been reported in wild *C. elegans* isolates (Hahnel *et al*. 2018), and of these S145F and M257I can be induced by EMS. These alleles may differ from the BZ resistant mutations in the species described above in that they need not retain wild-type functions.

Another indication that a limited number of missense changes in *ben-1* can mutate to ABZ resistance is that we found two pairs of mutations that cause different amino acid changes in the same codon. *e1880* (G104D) and *sb145* (G104S) both alter amino acid 104 while *sb144* (G142E) and *sb164* (G142R) are both at position 142. This overlap could indicate that these positions are unique in that could be critical for protein function.

Our screen appeared to efficiently identify nonsense and splicing mutations more than missense alleles, again implying an unexpected rarity of amino acid changes that can confer resistance. We isolated four of 35 possible EMS-induced *ben-1* nonsense and splicing mutations. By contrast, we found only 12 of the 389 possible *ben-1* missense mutations that can be induced by EMS (excluding the non-conserved C-terminus, K.M. Tahsin Hassan Rahit and M. Tarailo-Graovac, personal communication). Perhaps a more compelling example of the paucity of missense alleles that can confer ABZ resistance is that in wild *C. elegans* populations, where *ben-1* mutations may arise after environmental BZ exposure, Hahnel *et al*. (2018) found that only three of 25 resistance mutations were missense.

Thus, the assumption that high conservation of β-tubulin indicates that most missense alleles would result in severe loss of function may not be valid. This hypothesis is consistent with a systematic survey of the *S. cerevisiae* β-tubulin gene. Reijo *et al*. (1994) changed clusters of charge amino acids to alanine and found that only 11of 55 alleles were lethal (although many viable alleles would cause fitness costs in nature). Only five were strongly resistant to benomyl. This result suggests that in *C. elegans* most *ben-1* missense mutations would not confer ABZ resistance as they could retain sufficient function to deliver the BZ poison to the microtubule.

### *ben-1* is the major ABZ target in *C. elegans*

With the widespread use of ABZ in human populations, it is critical to understand the genetics of nematode drug resistance. *ben-1* is clearly the major target in *C. elegans* under laboratory conditions. The resistant allele *sb163* (A63V) that retains *ben-1(+)* function could represent a new mutation that may arise in parasites. If most *ben-1* missense alleles do not confer resistance, parasitic nematodes species with a redundant Y200 containing paralogs might be more likely to acquire resistance through nonsense and deletion mutations than the F167Y, E198A, and F200Y commonly found in BZ resistant parasites. Additional genes also influence BZ resistance in both wild *C. elegans* and in parasitic nematodes (Hahnel *et al*. 2018; Zamanian *et al*. 2018; Furtado *et al*. 2019a). We did find one mutation (*sb156*) with no lesions in the *ben-1* coding region and this mutant had the weakest resistance in our study. Further optimization of our screens to recover weak resistance may uncover additional genes.

## FUNDING

This work was supported by grants from the Canadian Institute of Health Research (CIHR) and the Natural Science and Engineering Council (NSERC) of Canada to P.E.M. C.M.K, J.S.G., and E.C.A. were funded by NIH R01 AI153088. Some strains were provided by the CGC, which is funded by NIH Office of Research Infrastructure Programs (P40 OD010440).

## COMPETING INTERESTS

None

## ACKNOWLEDGMENTS

We thank the members of the labs of D. Hansen, J.D. McGhee and M. Tarailo-Graovac for comments throughout this project. We also thank WormBase. F. Jean, S. Stasiuk, and F. Snider are thanked for technical support. K.M. Tahsin Hassan Rahit and M. Tarailo-Graovac provided calculation for mutation frequencies. L.M.P. was part of the Waterloo undergraduate Co-op program.

---

**Supplemental Figure 1.**
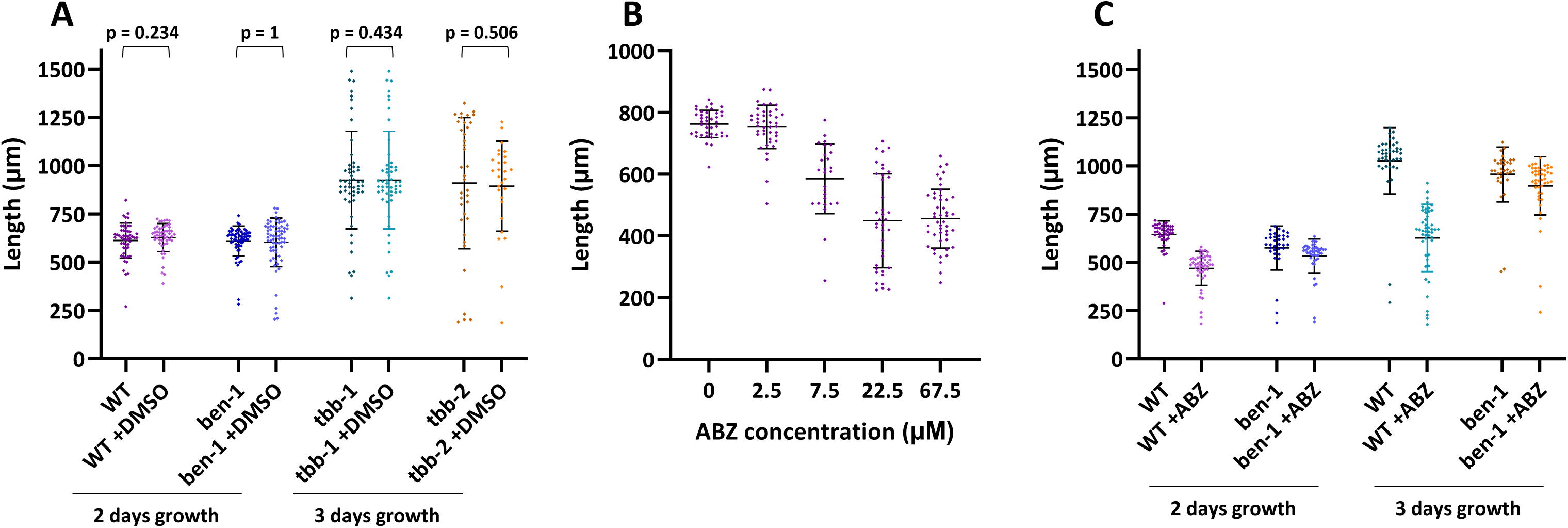
Optimization of growth assays. (A) ABZ was added to plates dissolved in DMSO. 0.5% DMSO did not affect growth of wild type *ben-1, tbb-1* or *tbb-2* and so was not included in control plates. (B) Dose response curves of wild type grown at 20° and measured after 1 or 2 days growth. 7.5 μM was chosen so that both increased and decreased ABZ sensitivity could be assessed. (C) Comparisons of growth differences between wild type and *ben-1* null alleles on 7.5 μM ABZ after 2 or 3 days of growth. Differences were maximized at 3 days, which was before the next generation begins to hatch. This allowed scoring of arrested animals. Two-tailed Mann-Whitney Rank Sum test were used to calculate p values. Mean and standard deviations are indicated.

**Supplemental Figure 2.**
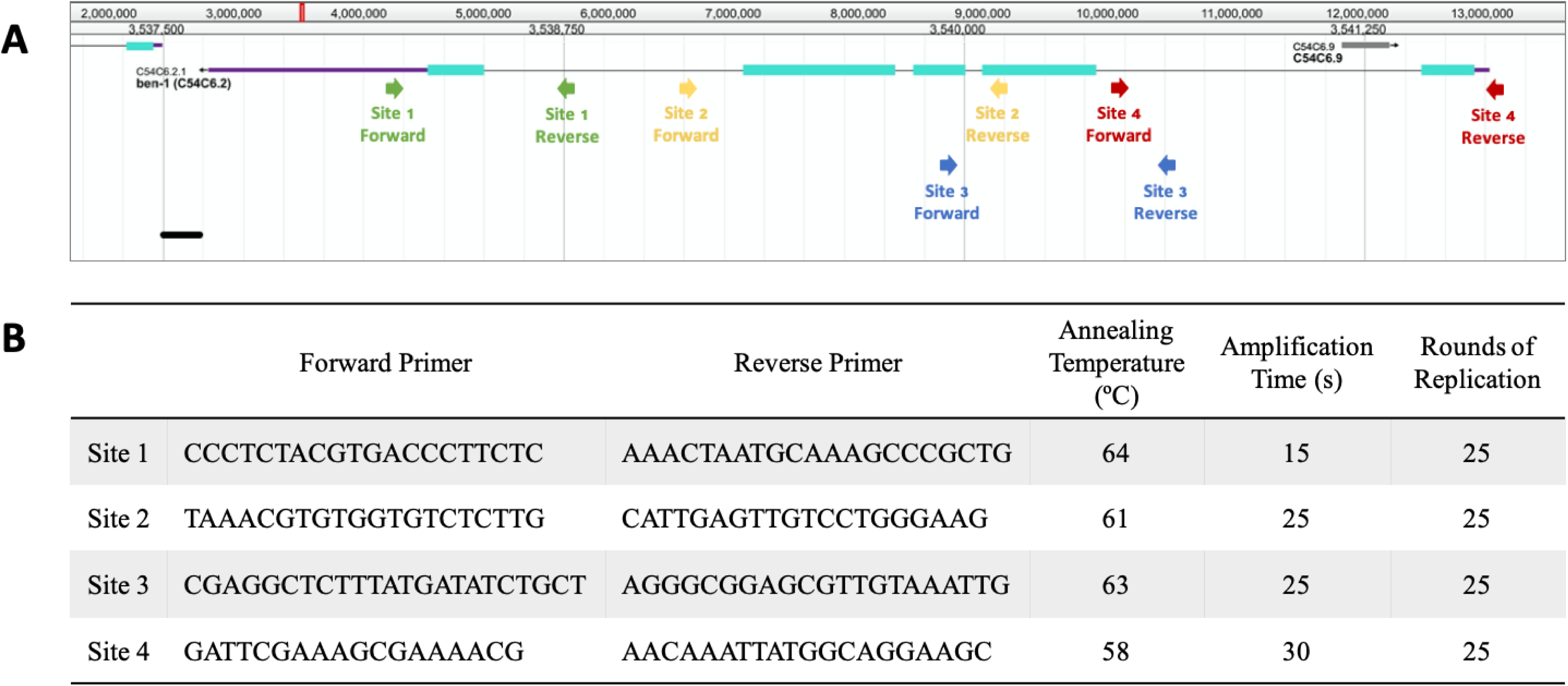
(A) Position of primers used for *ben-1* genomic sequencing in a WormBase screenshot. Exons are indicated in blue. (B) Primer sequences and PCR conditions for *ben-1* sequencing.

**Supplemental Figure 3.**
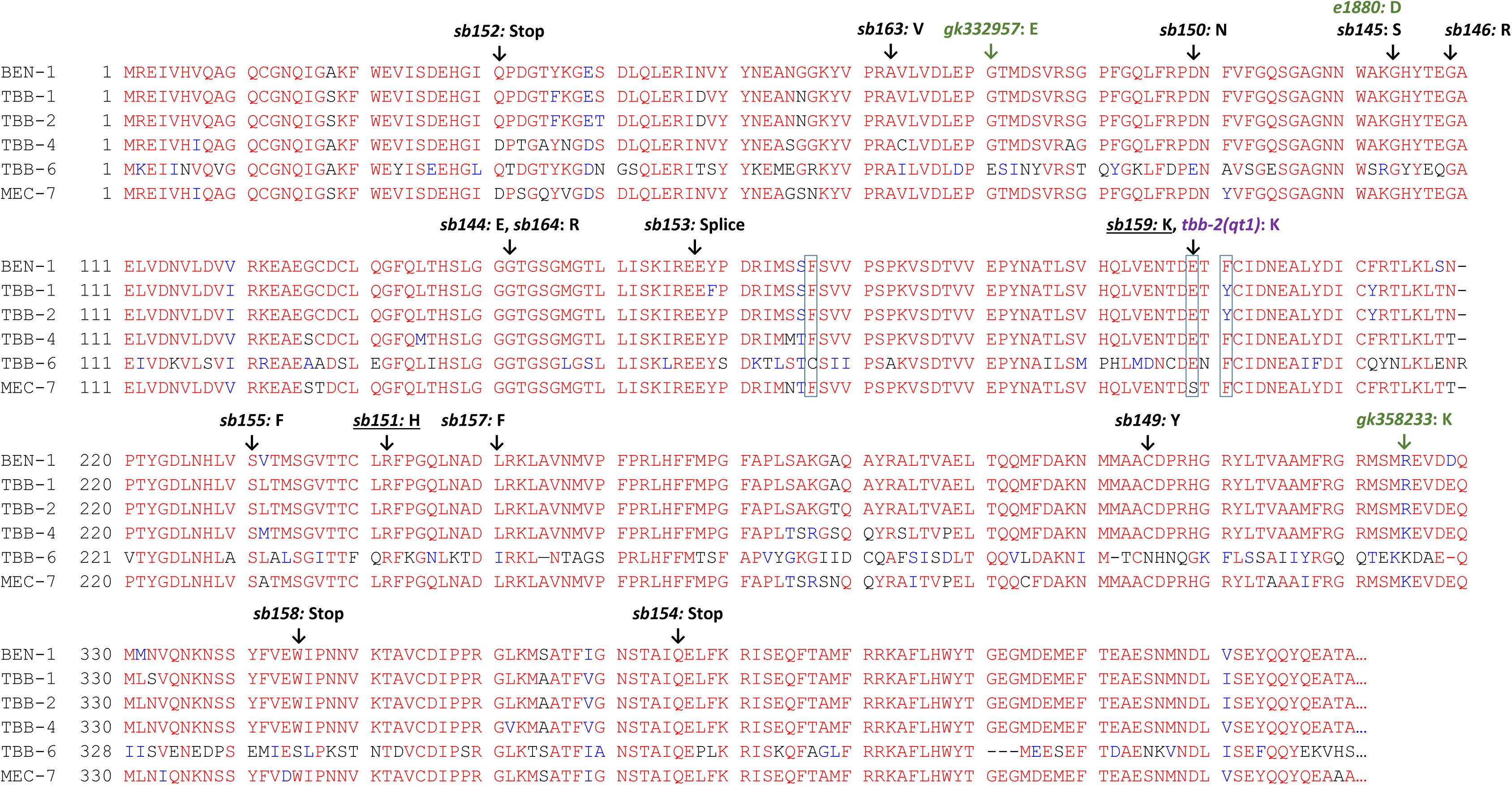
Alignment of BEN-1 with the other *C. elegans* β-tubulins. Positions of the ABZ mutants that we isolated are shown in black along with the canonical allele *e1880*. Boxed residues are frequently mutated in parasitic nematodes. Underlined alleles are found in BZ resistant mutants of other organisms. Green indicates alleles found in the Million Mutant Project and purple denotes the change in *tbb-2(qt1).* Note that numbering is altered for TBB-6 due to an insertion at 219 relative to other tubulins. Sequences are truncated to exclude the non-conserved C-terminal regions.

**Supplemental Table 1.**
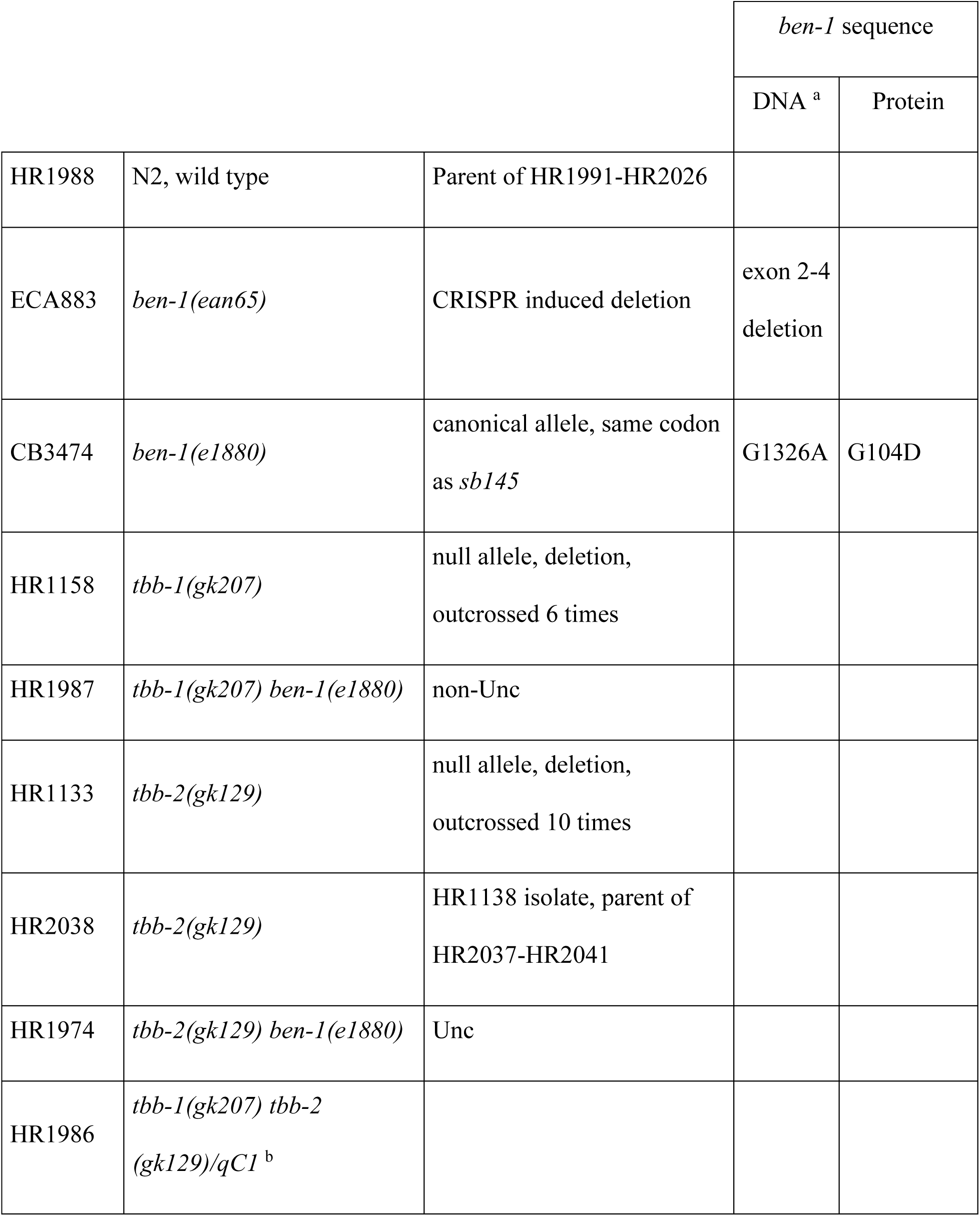
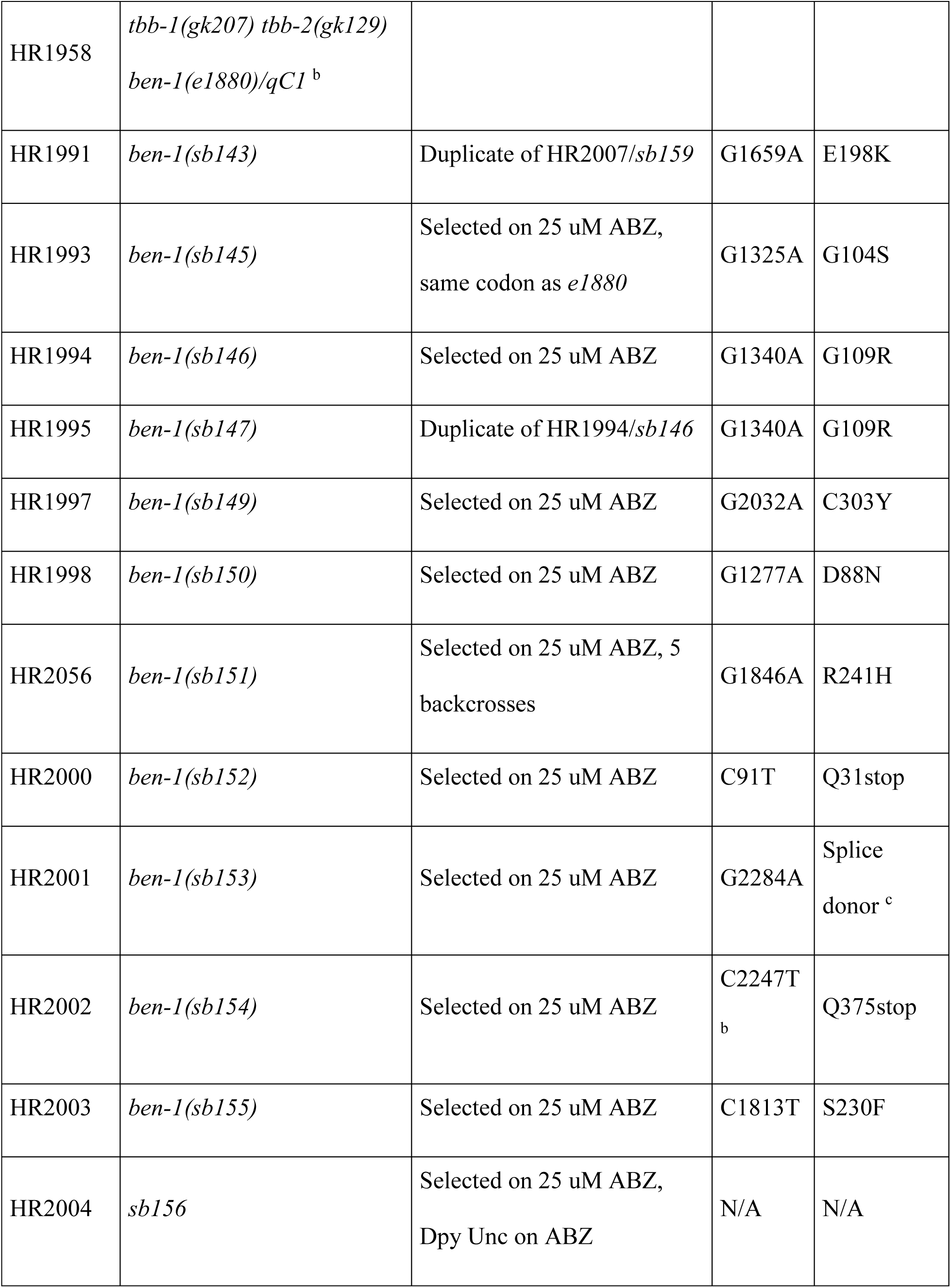
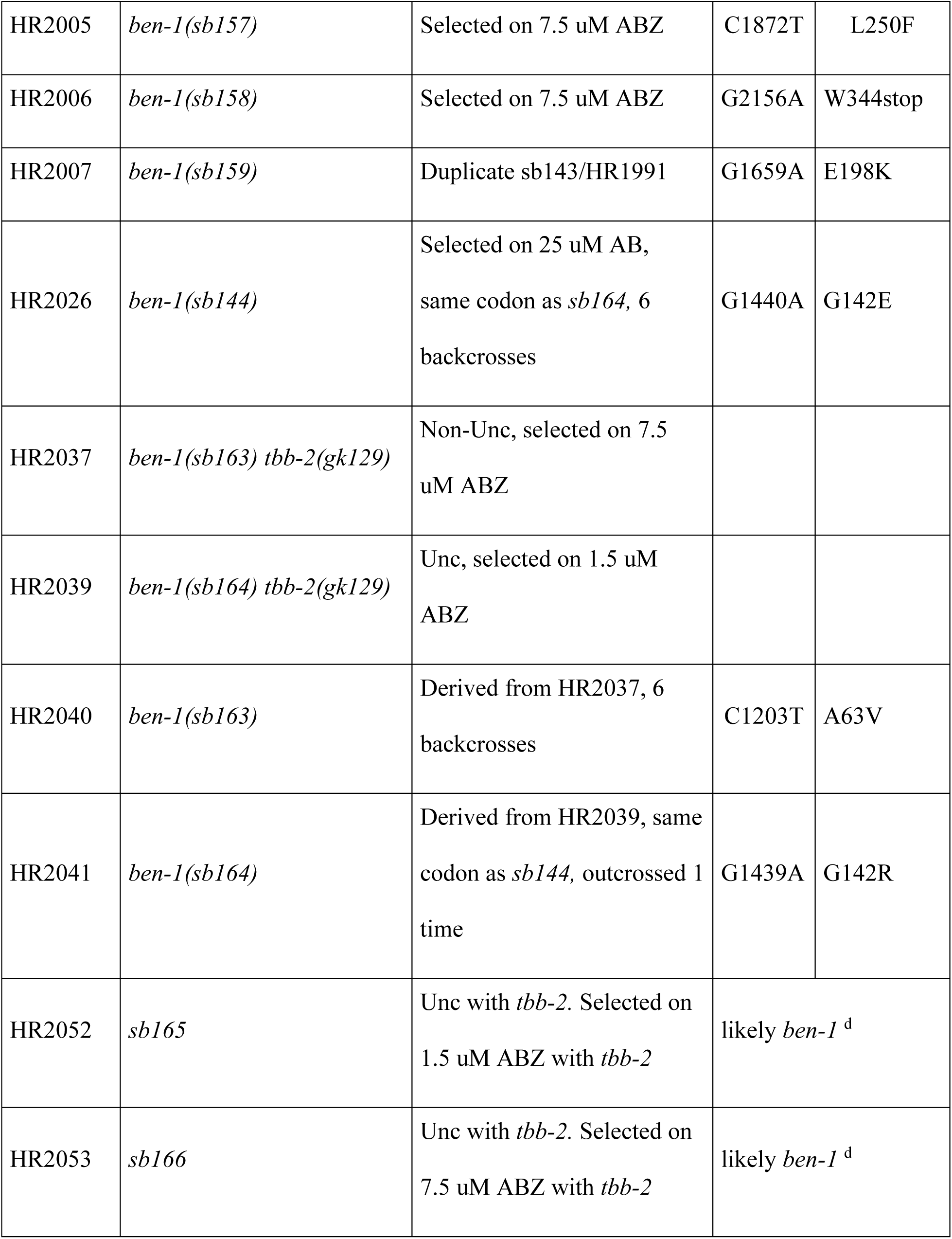
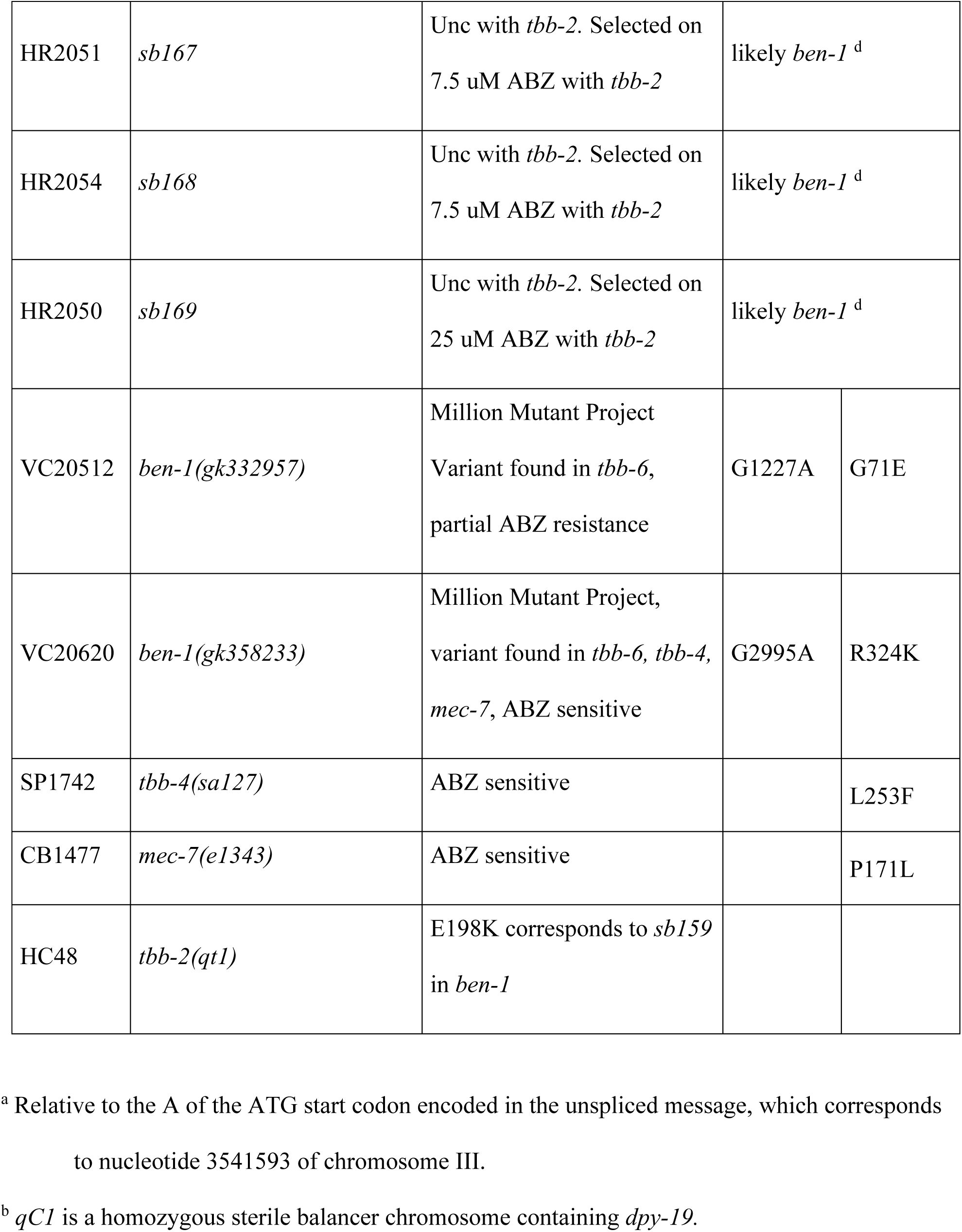
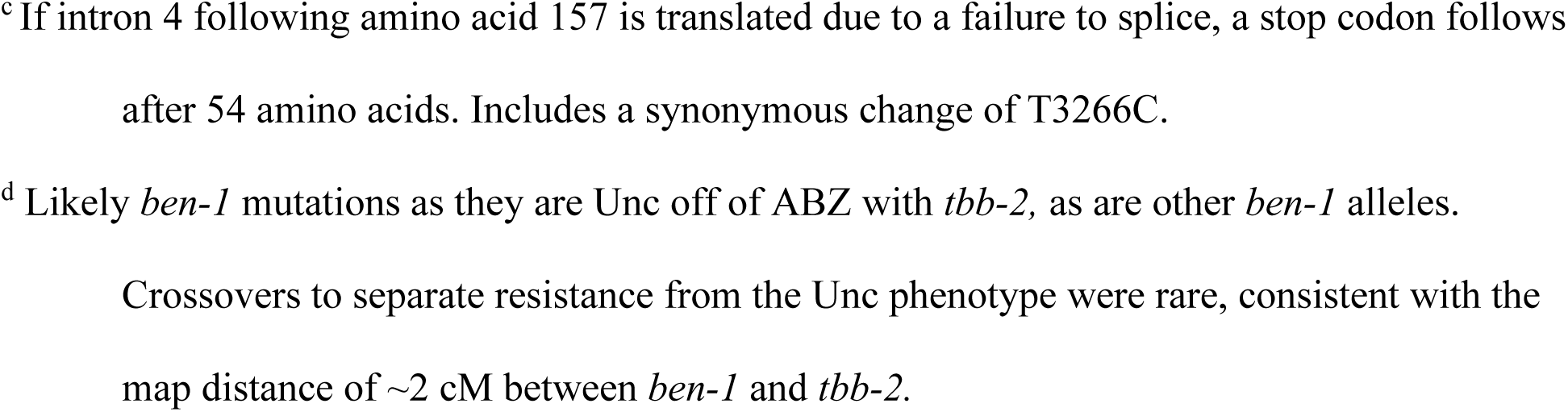
Strain genotypes

|  |  |  | *ben-1* sequence |  |
| --- | --- | --- | --- | --- |
|  |  |  | DNA <sup>a</sup> | Protein |
| HR1988 | N2, wild type | Parent of HR1991-HR2026 |  |  |
| ECA883 | *ben-1(ean65)* | CRISPR induced deletion | exon 2-4<br>deletion |  |
| CB3474 | *ben-1(e1880)* | canonical allele, same codon<br>as *sb145* | G1326A | G104D |
| HR1158 | *tbb-1(gk207)* | null allele, deletion,<br>outcrossed 6 times |  |  |
| HR1987 | *tbb-1(gk207) ben-1(e1880)* | non-Unc |  |  |
| HR1133 | *tbb-2(gk129)* | null allele, deletion,<br>outcrossed 10 times |  |  |
| HR2038 | *tbb-2(gk129)* | HR1138 isolate, parent of<br>HR2037-HR2041 |  |  |
| HR1974 | *tbb-2(gk129) ben-1(e1880)* | Unc |  |  |
| HR1986 | *tbb-1(gk207) tbb-2*<br>*(gk129)/qC1 <sup>b</sup>* |  |  |  |
| HR1958 | *tbb-1(gk207) tbb-2(gk129)*<br>*ben-1(e1880)/qC1<sup>b</sup>* |  |  |  |
| HR1991 | *ben-1(sb143)* | Duplicate of HR2007/*sb159* | G1659A | E198K |
| HR1993 | *ben-1(sb145)* | Selected on 25 uM ABZ,<br>same codon as *e1880* | G1325A | G104S |
| HR1994 | *ben-1(sb146)* | Selected on 25 uM ABZ | G1340A | G109R |
| HR1995 | *ben-1(sb147)* | Duplicate of HR1994/*sb146* | G1340A | G109R |
| HR1997 | *ben-1(sb149)* | Selected on 25 uM ABZ | G2032A | C303Y |
| HR1998 | *ben-1(sb150)* | Selected on 25 uM ABZ | G1277A | D88N |
| HR2056 | *ben-1(sb151)* | Selected on 25 uM ABZ, 5<br>backcrosses | G1846A | R241H |
| HR2000 | *ben-1(sb152)* | Selected on 25 uM ABZ | C91T | Q31stop |
| HR2001 | *ben-1(sb153)* | Selected on 25 uM ABZ | G2284A | Splice<br>donor<sup>c</sup> |
| HR2002 | *ben-1(sb154)* | Selected on 25 uM ABZ | C2247T<sup>b</sup> | Q375stop |
| HR2003 | *ben-1(sb155)* | Selected on 25 uM ABZ | C1813T | S230F |
| HR2004 | *sb156* | Selected on 25 uM ABZ,<br>Dpy Unc on ABZ | N/A | N/A |
| HR2005 | *ben-1(sb157)* | Selected on 7.5 uM ABZ | C1872T | L250F |
| HR2006 | *ben-1(sb158)* | Selected on 7.5 uM ABZ | G2156A | W344stop |
| HR2007 | *ben-1(sb159)* | Duplicate sb143/HR1991 | G1659A | E198K |
| HR2026 | *ben-1(sb144)* | Selected on 25 uM AB,<br>same codon as *sb164*, 6<br>backcrosses | G1440A | G142E |
| HR2037 | *ben-1(sb163) tbb-2(gk129)* | Non-Unc, selected on 7.5<br>uM ABZ |  |  |
| HR2039 | *ben-1(sb164) tbb-2(gk129)* | Unc, selected on 1.5 uM<br>ABZ |  |  |
| HR2040 | *ben-1(sb163)* | Derived from HR2037, 6<br>backcrosses | C1203T | A63V |
| HR2041 | *ben-1(sb164)* | Derived from HR2039, same<br>codon as *sb144*, outcrossed 1<br>time | G1439A | G142R |
| HR2052 | *sb165* | Unc with *tbb-2*. Selected on<br>1.5 uM ABZ with *tbb-2* | likely *ben-1*<sup>d</sup> |  |
| HR2053 | *sb166* | Unc with *tbb-2*. Selected on<br>7.5 uM ABZ with *tbb-2* | likely *ben-1*<sup>d</sup> |  |
| HR2051 | *sb167* | Unc with *tbb-2*. Selected on 7.5 uM ABZ with *tbb-2* | likely *ben-1*<sup>d</sup> |  |
| HR2054 | *sb168* | Unc with *tbb-2*. Selected on 7.5 uM ABZ with *tbb-2* | likely *ben-1*<sup>d</sup> |  |
| HR2050 | *sb169* | Unc with *tbb-2*. Selected on 25 uM ABZ with *tbb-2* | likely *ben-1*<sup>d</sup> |  |
| VC20512 | *ben-1(gk332957)* | Million Mutant Project<br>Variant found in *tbb-6*,<br>partial ABZ resistance | G1227A | G71E |
| VC20620 | *ben-1(gk358233)* | Million Mutant Project,<br>variant found in *tbb-6*, *tbb-4*,<br>*mec-7*, ABZ sensitive | G2995A | R324K |
| SP1742 | *tbb-4(sa127)* | ABZ sensitive |  | L253F |
| CB1477 | *mec-7(e1343)* | ABZ sensitive |  | P171L |
| HC48 | *tbb-2(qt1)* | E198K corresponds to *sb159*<br>in *ben-1* |  |  |

<sup>a</sup> Relative to the A of the ATG start codon encoded in the unspliced message, which corresponds to nucleotide 3541593 of chromosome III.
<sup>b</sup> *qC1* is a homozygous sterile balancer chromosome containing *dpy-19*.
<sup>c</sup> If intron 4 following amino acid 157 is translated due to a failure to splice, a stop codon follows after 54 amino acids. Includes a synonymous change of T3266C.
<sup>d</sup> Likely *ben-1* mutations as they are Unc off of ABZ with *tbb-2*, as are other *ben-1* alleles.
Crossovers to separate resistance from the Unc phenotype were rare, consistent with the map distance of ~2 cM between *ben-1* and *tbb-2*.

**Supplemental Table 2.** Mutant screens

| Parent | ABZ (μM) | Gravid F1 | Number of stains |
| --- | --- | --- | --- |
| Wild-type | 7.5 | 140<sup>a</sup> | 1 |
|  | 7.5 | 410<sup>b</sup> | 1 |
|  | 25 | 3,960<sup>a</sup> | 12<sup>c</sup> |
|  | 50 | 240<sup>a</sup> |  |
|  | haploid genomes | 9,500 |  |
| *ben-1* mutations/gamete |  | 1/730 |  |
| *tbb-2* | 1.5 | 1,275 | 1 |
|  | 7.5 | 3,775 | 1 |
|  | haploid genomes | 10,100 |  |
| *ben-1* mutations/gamete |  | 1/5050 |  |
| *tbb-2* | 1.5 | 12,280 | 1<sup>d</sup> |
|  | 3 | 2,060 |  |
|  | 7.5 | 3,620 | 3<sup>d</sup> |
|  | 25 | 6,470 | 1<sup>d</sup> |
|  | 50 | 1,350 |  |
|  | haploid genomes | 51,560 |  |
|  | *ben-1* mutations/gamete<sup>c</sup> | 1/10310 |  |

\* Gravid F1 were grown from mutagenized worms in the absence of drug and 20-30 were picked to each ABZ plate. In all other cases, mutagenized worms were directly plated on ABZ and gravid F1's present a week later were counted from a sampling of plates.
<sup>b</sup> Mutagenized animals were picked directly to ABZ plates.
<sup>c</sup> Excludes *sb156*, which has no coding changes in *ben-1*.
<sup>d</sup> Likely *ben-1* mutations as they are Unc off of ABZ with *tbb-2* as are other *ben-1* mutations. Crossovers to separate resistance from the Unc phenotype were rare, consistent with the map distance of ~2 cM between *ben-1* and *tbb-2*.

